# Dividing subpopulation of *Escherichia coli* in stationary phase

**DOI:** 10.1101/614974

**Authors:** Arvi Jõers, Eliis Liske, Tanel Tenson

## Abstract

The bacterial growth cycle contains different phases: after the growth substrate is exhausted of the toxic waste products accumulate the growth stops. In this non-growing culture the number of colony forming bacteria remains constant or starts to decrease. It has been shown that during prolonged incubation there is constant growth and death of bacteria and certain mutant populations take over the culture. Here we show that the dynamic cell division and death balance can be obtained even before mutants take over the culture.

## Introduction

The bacterial growth curve is classically divided into four steps: lag phase, exponential growth phase, stationary phase and death phase. Until the late 1980s and early 1990s the exponential growth phase was considered as the major characteristic of a bacterium isolate and was a main target of investigations. Later research efforts have been increasingly focusing also on other phases of the growth cycle, as in nature exponential growth covers only a small period of the ecological life cycle of a bacterium [1].

Stationary phase occurs when bacteria have either exhausted some necessary nutrient from the medium or waste products have accumulated to the level that prohibits further growth [1,2]. The number of bacteria, usually measured by colony forming units (CFU) or optical density (OD), reaches a plateau and stay unchanged for some time. This stable situation at the bulk level can arise from two different scenarios at the individual cell level. All the cells may be indeed in nondividing state and truly static. Alternatively, constant cell death and division takes place in such a manner that the total number of (alive) cells stays the same. While the latter scenario is frequently speculated about there is surprisingly few direct studies on the matter that do not involve selection for mutants.

After a prolonged incubation in stationary phase mutants arise that take over the culture [3]. These are called growth advantage in stationary phase (GASP) mutants and usually have mutations in *rpoS, lrp* and few other genes [4]. In *Escherichia coli* these mutants are studied in LB medium, where they appear in 10 days [3].

We have previously observed that all cells stop dividing when entering the stationary phase in LB medium and stay homogenously nondividing during the first day(s) [5]. In this paper we analyze at the single cell level what happens between the early stationary phase and the appearance of GASP mutants.

## Results

### Dividing subpopulation appears after a few days in stationary phase

During the measurement of colony forming units (CFU) in stationary phase we noticed that they fluctuate (data not shown). After initial decline their level increases again after 3 - 5 days in LB. To investigate this phenomenon further we employed 2-color flow cytometric method for following cell division at the single cell level [6]. Briefly, cells contain two plasmids: one carrying inducible GFP gene and the other inducible Crimson gene. Cells are grown with Crimson expression induced and after reaching stationary phase this inducer is removed. The second inducer for GFP is added to the stationary phase culture, but because cells have ceased their activities, GFP is not expressed. However, if some cells become active again they will become GFP positive and will lose their Crimson content because of dilution by cell division.

When we applied this analysis on stationary phase culture in LB the dividing subpopulation appeared in 4-day culture (Figure 1). During the first 3 days no changes can be detected and culture remains static. On the 4th day however the GFP positive cells appear indicating that some cells have started protein synthesis. Majority of GFP positive cells have also become Crimson negative, indicating that they have divided several times. Nevertheless, there is a significant GFP positive, Crimson positive subpopulation, containing cells that have become active but have not divided yet. This allows us to estimate the frequency of cells reactivating their metabolism. Based on flow cytometry data we calculate it to be 3.4 +− 2.6 % (average +− standard deviation). This is an underestimation, as some active cells have divided and became Crimson negative. Despite of that this value by far exceeds any reported mutation frequencies in *E. coli*, suggesting that at least the majority of active subpopulation is not caused by specific mutation.

**Figure 1.**
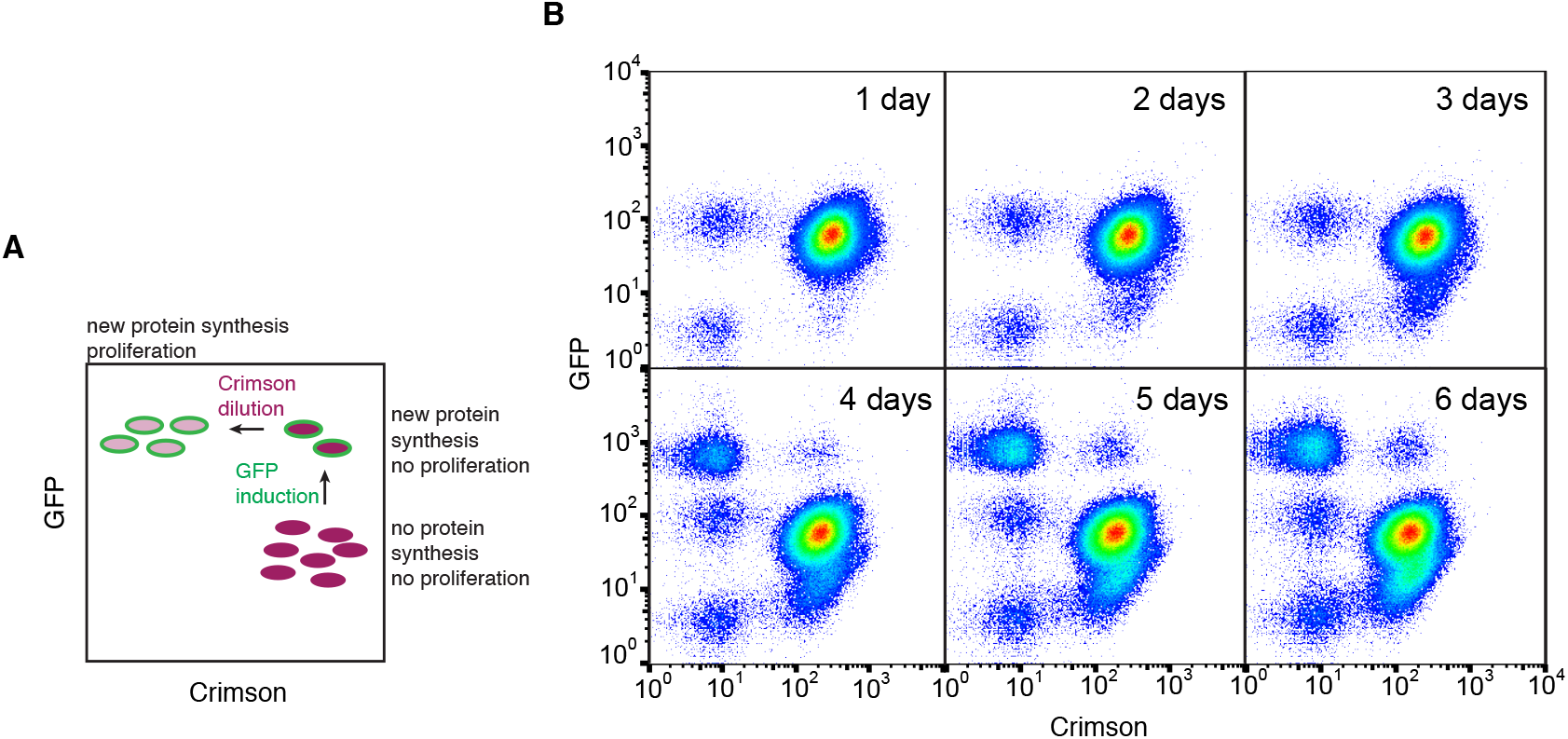
Dividing cells appear in 4-day culture. **A.** Schematic presentation of different subpopulations in stationary phase. **B.** Subpopulations in stationary phase. Cells were grown in LB with Crimson induced and after reaching stationary phase the Crimson inducer was removed and GFP inducer added (1-day culture). After 4 days GFP positive population emerges.

### Eliminating cell division prevents increase in CFU numbers

To test if the increase of CFU in stationary phase is indeed caused by *de novo* cell division we used ampicillin to eliminate dividing cells. Ampicillin is able to lyse growing and dividing cells by disrupting their cell wall, but is harmless to nondividing cells. When ampicillin was added to stationary phase culture it did not affect the CFU count during the first 3 days, but prevented both the increase of CFU at the day 4 (Figure 2A) and the appearance of GFP positive subpopulation at the same time (Figure 2B). This indicates that *de novo* cell division is the reason behind the increase of CFU.

**Figure 2.**
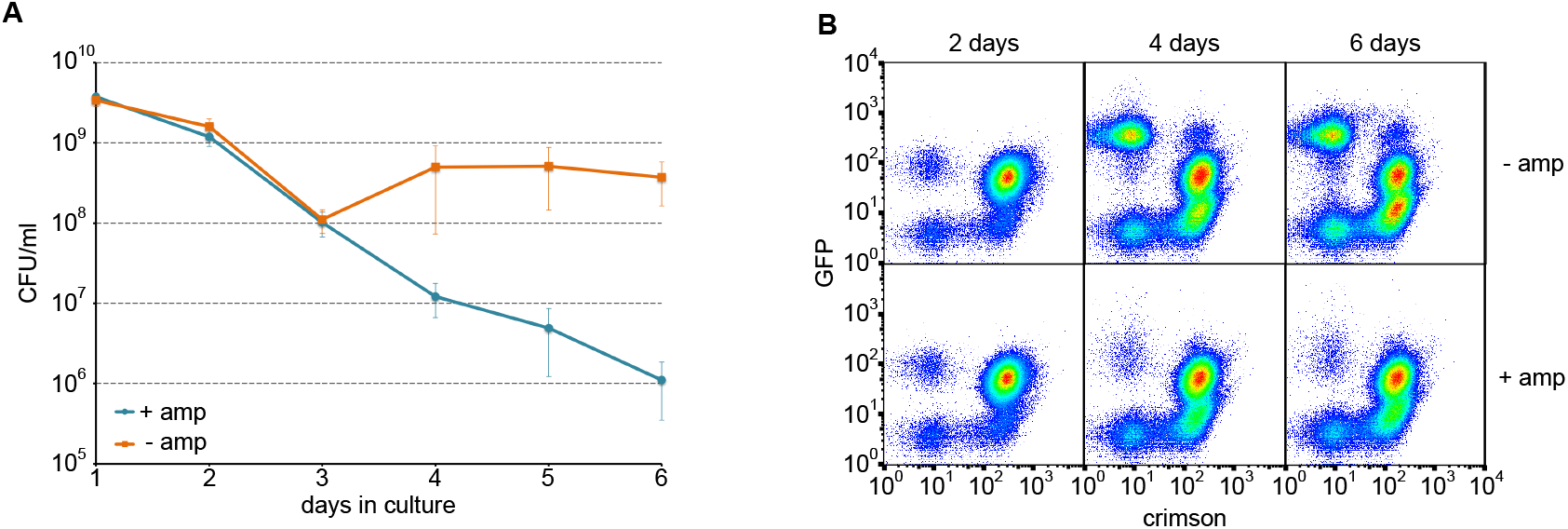
Ampicillin (amp) prevents the appearance of active population. Cells were incubated in LB stationary phase culture for 6 days with or without amp. Both viability test (A) and flow cytometry analysis (B) indicate the inability of active subpopulation to emerge in the presence of amp.

### Dividing cells do not have a phenotype of a GASP mutant

GASP mutants, by definition, are able to outcompete wt strain during the stationary phase. Once isolated, the phenotype of these mutants is stable and they can take over the wt culture in competition experiments. We wanted to test if cells from the dividing subpopulation have acquired some mutation that allows them to outcompete wt strain. To do that we first isolated naturally occurring rifampicin (Rif) or nalidixic acid (Nal) resistant mutants and cultivated them in LB for 5 days. Both displayed phenotypic heterogeneity with dormant, active, and dividing cells present (data not shown). We then sorted dividing and dormant cells in cell sorter and plated them on agar plates to obtain single colonies (Figure 3A).

**Figure 3.**
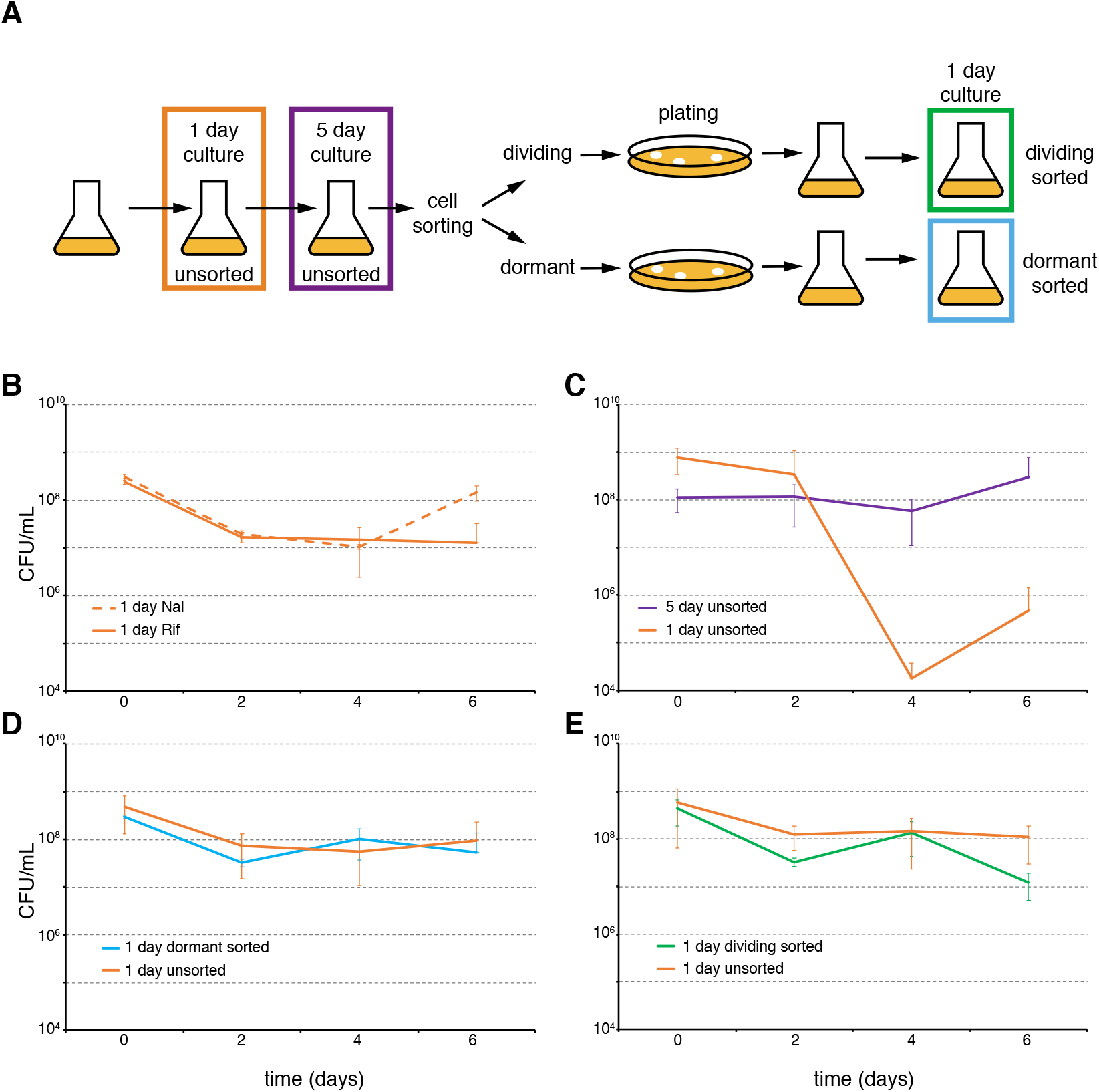
Dividing cells do not have GASP phenotype. **A.** Experimental scheme to obtain 1 day cultures with different histories. **B.** When mixed together, unsorted 1 day old cultures are neutral to each other. The averages and standard errors of 3 independent experiments are shown. **C.** 5 day unsorted culture outcompetes 1 day unsorted culture. The averages and standard errors of 6 independent experiments are shown. **D.** Sorted cells from dormant population do not outcompete 1 day unsorted culture. The averages and standard errors of 4 independent experiments (colonies) are shown. **E.** Sorted cells from dividing population do not outcompete 1 day unsorted culture. The averages and standard errors of 4 independent experiments (colonies) are shown.

Both sorted and unsorted strains were grown into stationary phase in LB and cultures with different resistance markers (Rif or Nal) were mixed together. 1 day old unsorted cultures do not gain significant growth advantage over one another in mixed culture, demonstrating marker neutrality (Figure 3B). Unsorted 5 day culture is able to outcompete unsorted 1 day culture if incubated together for more than 4 days (Figure 3C). When unsorted and sorted 1 day cultures were mixed together, neither of them was able to outcompete the other (Figure 3D and 3E). It did not matter that dividing sorted cells were able to grow in previous stationary phase. After sorting and regrowth this phenotype had disappeared and they were like unsorted cells. This suggests that cell division in stationary phase is initially caused by transient phenotypic change that is not based on GASP mutation. This phenotypic change is behind the heterogeneity in the culture and also enables the 5 day culture to outcompete 1 day culture (Figure 3C).

## Discussion

*E. coli* long term stationary phase culture in rich medium is known to be dynamic. The appearance of GASP (growth advantage in stationary phase) mutants was described more than 25 years ago and several GASP mutations have been identified [3]. It takes 10 days to develop detectable GASP mutants in LB. Here we describe that this is preceded by the appearance of dividing subpopulation of cells. These arise far too frequently to be mutants by themselves, but probably form a prerequisite for GASP mutation to occur.

Cell division requires growth substrate, so there must be one present in 4 day stationary phase. Two possible sources can be envisioned. At first cells might secrete a secondary metabolite during the initial growth phase that can be used as a carbon source later. Alternatively, cells may die during the stationary phase, leak out their content and provide carbon source for surviving population. The second explanation is in accord with the fact that the CFU drops close to two orders of magnitude during the first 3 days in stationary phase indicating that most of the cells lose their viability.

Our results demonstrate that, depending on conditions, stationary phase in *E. coli* can be either static or dynamic. In the latter case phenotypic heterogeneity precedes the mutational one. Similar dynamics have been described also for antibiotic resistance mutations. There dormant subpopulation survives the periodic antibiotic treatment and gives rise to growing subpopulation during the antibiotic-free intermediate periods [7]. This facilitates the appearance of resistant mutants, providing a window of opportunity for these mutations to occur. In our case growing subpopulation in stationary phase provides more chances for GASP mutations to occur and eventually they take over the culture. Only a small number of cells start growing after 4 days in stationary phase. This might reflect the loss of viability in majority of population, although there are live cells among GFP-negative population. This could also be a manifestation of bet-hedging strategy, creating subpopulations with different phenotypes, optimal for different future environments [8]. We have shown before that heterogeneous growth resumption ensures population survival in the case of heat shock [6]. We cannot say if growth in stationary phase is a true bet-hedging behavior, but it does seem to promote a long-term survival of the microbial population.

## Methods

### Bacterial strains and media

E. coli strain BW25113 (F-, Δ(araD-araB)567, ΔlacZ4787(::rrnB-3), λ-, rph-1, Δ(rhaD-rhaB)568, hsdR514) was used throughout the study. Plasmids pET-GFP and pBAD-Crimson [6] were used to express GFP and E2-Crimson respectively.

### Detecting growth in stationary phase

BW25113 cells containing pET-GFP and pBAD-Crimson plasmids were grown in LB supplemented with kanamycin (25μg/mL) and chloramphenicol (25μg/mL) for plasmid retention, and arabinose (1 mM) for Crimson induction. In parallel the same strain was grown without arabinose. On the next day both cultures were centrifuged and the supernatant from arabinose-containing culture was replaced with sterile-filtered supernatant from cells grown without arabinose. This removes crimson inducer while retaining the stationary phase conditions. IPTG (1 mM) was added to induce GFP expression in cells capable of protein synthesis. Samples for flow cytometry were taken every day for 6 days.

LB (Difco) was prepared freshly and sterilized by filtration through 0.22 μm filter. We noticed that the exact time of the appearance of dividing subpopulation was a little different for different LB medium batches. With the batch used in this paper active cells appear at the 4th or 5th day, whereas with some earlier batches dividing cells were visible already at the 3rd day. Regardless of the exact timing the overall phenomenon was the same every time.

### Flow cytometry

Samples for flow cytometry were taken daily, mixed 1:1 with sterile filtered 30% glycerol in PBS and stored at −70° C pending analysis. Flow cytometry was carried out using LSRII (BD Biosciences) equipped with blue (488 nm) and red (638 nm) lasers. The detection windows for GFP and Crimson were 530±15 nm and 660±10 nm respectively. Flow cytometry samples stored at −70° C were thawn at room temperature and 5 μl of cell suspension was diluted in 0.5 ml PBS to prepare for flow cytometer. At least 20 000 events were analyzed for every sample.

Cell sorting was done using FACSAria (BD Biosciences) with the same detection parameters using 70 μm nozzle. GFP-negative/Crimson-positive (dormant) and GFP-positive/Crimson-negative (growing) cells were sorted into sterile PBS and plated onto LB plates to obtain individual colonies.

### Competition experiments

Two competing strains (one Rif and the other Nal resistant) were mixed together, 1 ml of each, and incubated at the shaker in 37° C throughout the experiment. Every day a 10 μL sample was taken and used to prepare serial dilutions on 96-well microtiter plate in sterile PBS. 5 μL from every dilution was spot-plated on LB plates containing either Rif (100 μg/mL) or Nal (20 μg/mL) and the colonies were counted on the next day.

## Acknowledgements

The work was supported by European Union from the European Regional Development Fund through the Centre of Excellence in Molecular Cell Engineering (2014-2020.4.01.15-0013) and grant from the Estonian Research Council (PRG335).

